# Changes in cross-frequency coupling following closed-loop auditory stimulation in non-rapid eye movement sleep

**DOI:** 10.1101/810861

**Authors:** Krugliakova Elena, Volk Carina, Jaramillo Valeria, Sousouri Georgia, Huber Reto

## Abstract

The activity of different brain networks in non-rapid eye movement (NREM) sleep is regulated locally in an experience-dependent manner, reflecting the extent of the network load during wakefulness. In particular, improved task performance after sleep correlates with the local post-learning power increase of neocortical slow waves and faster oscillations such as sleep spindles and their temporal coupling. Recently, it was demonstrated that by targeting slow waves in a particular region at a particular phase with closed-loop auditory stimulation it is possible to locally manipulate slow-wave activity and interact with training-induced neuroplastic changes. Based on this finding, we tested whether closed-loop auditory stimulation targeting the up-phase of slow-waves over the right sensorimotor area might affect power in delta, theta and sigma bands and coupling between these oscillations within the circumscribed region. We demonstrate that while closed-loop auditory stimulation globally enhances power in delta, theta and sigma bands, changes in cross-frequency coupling of these oscillations were more spatially restricted. In particular, stimulation induced a significant decrease of delta-theta coupling in frontal channels, within the area of the strongest baseline coupling between these frequency bands. In contrast, a significant increase in delta-sigma coupling was observed over the right parietal area, located directly posterior to the target electrode. These findings suggest that closed-loop auditory stimulation locally modulates coupling between delta phase and sigma power in a targeted region, which could be used to manipulate sleep-dependent memory formation within the brain network of interest.

## 1. Introduction

A substantial body of evidence highlights the importance of non-rapid eye movement (NREM) sleep for experience-dependent neuroplasticity and associated improvements in behavioral performance after a bout of sleep (Rasch and Born, 2013; Tononi and Cirelli, 2014). NREM sleep electroencephalographic (EEG) recordings are hallmarked by neocortical slow waves (1-4 Hz) and sleep spindles, the waxing and waning activity in the sigma frequency band (10-16 Hz). It is well established that slow waves orchestrate faster oscillations, with the up-phase of slow waves being associated with increased sigma band activity, and sigma oscillations, in turn, group gamma activity in their troughs (Mölle et al., 2002; Staresina et al., 2015). This precise cross-frequency phase-amplitude coupling is thought to organize communication across multiple brain areas and to facilitate coordinated information processing in brain networks during NREM sleep (Cox et al., 2014a).

Not only the power and spatial distribution of slow waves and sigma activity (Gais et al., 2002; Huber et al., 2004a; Mander et al., 2013, 2014; Mölle et al., 2002; Niknazar et al., 2015), but especially their temporal coupling seems to contribute to sleep-dependent benefits of behavioral performance (Diekelmann and Born, 2010; Walker and Stickgold, 2006). Similar to phase-amplitude coupling between slow waves and sigma, modulation of theta band activity (5-8 Hz) by the phase of slow waves has been described in several studies. Both in surface EEG and in intracranial (iEEG) recordings, theta power was increased during the transition to the down phase of slow waves, with the last trough of theta-burst coinciding with the slow-wave down-phase (Cox et al., 2014a, 2019; Gonzalez et al., 2018; Klinzing et al., 2016).

Moreover, theta band activity in NREM sleep may play a role in sleep-dependent memory consolidation in humans. NREM theta oscillations were associated with the replay of the wake neuronal activity patterns during slow-wave sleep (Jiang et al., 2017; Schreiner et al., 2018). Furthermore, theta-range oscillations were linked to the benefits of targeted memory reactivation, a technique where learning-related cue stimuli (e.g. an odor or a sound) are presented during NREM sleep to promote sleep-dependent memory consolidation (Lehmann et al., 2016; Schreiner et al., 2015b, 2015a, 2018; Schreiner and Rasch, 2015). Thus, clear evidence exists both for coupling between slow waves and NREM theta, and for the association of theta band activity with memory replay. Nevertheless, theta is still not included as a primary component in the theoretical framework describing the functional importance of the temporal interaction between slow waves and faster rhythms for sleep-dependent neuroplasticity. A thorough description of the coupling between slow waves, theta, and spindles within the same data set might provide novel insights into the role of theta activity in these processes.

The manipulation of slow-wave activity during sleep by various techniques was instrumental in establishing causal relationships between sleep and learning-related processes (Fattinger et al., 2017; Del Felice et al., 2015; Landsness et al., 2009; Marshall et al., 2006; Prehn-Kristensen et al., 2014). However, whether cross-frequency coupling is critical for such causal relationships between sleep and learning is unknown. A first step would be to show that it is possible to non-invasively manipulate cross-frequency coupling. Closed-loop auditory stimulation, a novel approach to modulate slow waves, can be used to collect such evidence. It has been shown that these brief auditory stimuli, time-locked to the up phase of endogenous slow waves, not only induced trains of high-amplitude slow waves but also boosted sigma and theta activity (Garcia-Molina et al., 2018; Leminen et al., 2017; Ngo et al., 2013, 2015; Papalambros et al., 2017). The analyses of evoked event-related potentials (ERP) and event-related spectral perturbations (ERSP) show that they share the primary characteristics of spontaneously generated slow waves. Specifically, these evoked slow waves follow a similar time-frequency pattern, where the transition to a down-phase is associated with theta activity (∼ 0.4 s after stimulus onset) and the up-phase coincides with a peak of sigma activity (∼ 1 s after stimulus onset). Interestingly, this theta and sigma power enhancement coupled to evoked slow waves is not only found in studies using closed-loop auditory stimulation with pink noise bursts (Cox et al., 2014b; Leminen et al., 2017; Papalambros et al., 2017), but also in studies using targeted memory reactivation paradigms (Cairney et al., 2018; Lehmann et al., 2016). The functional relevance of this similarity between a specific reactivation of activity (by cueing) and the seemingly unspecific boost of slow waves by closed-loop auditory stimulation is unknown.

One important aspect to consider in this context is the local regulation of slow-wave activity. For example, sleep-dependent plasticity is reflected in locally enhanced slow-wave activity during sleep following intensive training on a particular task during wakefulness (Hanlon et al., 2012; Huber et al., 2004b). This observation suggests that the “need for sleep” increases as a function of learning-related brain plasticity. More recently, similar local effects of pre-sleep learning were also found to be related to cross-frequency coupling between slow waves and sigma activity (Cox et al., 2018; Helfrich et al., 2018; Yordanova et al., 2017). Hence, it might be that *local* enhancement of power and cross-frequency *interaction* of the main sleep EEG rhythms in the relevant brain regions is more crucial for the post-sleep performance gains as compared to *global* changes in these parameters. The critical question that arises is whether closed-loop auditory stimulation during sleep results in global or local effects.

Thus, the goal of our analysis was to test whether slow-waves, theta and sigma oscillations and their interactions can be modulated specifically in a particular brain region by closed-loop auditory stimulation. Spatially restricted modulation of cross-frequency coupling could be potentially used as a tool to interact with sleep-dependent memory formation within the brain network of interest.

## 2. Results

We analyzed high-density EEG recordings of 9 subjects (18 full-night recordings). Subjects participated in two-night sessions separated by 1 week: non-stimulation (SHAM) and closed-loop auditory stimulation in the up-phase (STIM) of real-time detected slow waves. Real-time closed-loop slow-wave detection during sleep stages N2 and N3 was performed as described in Fattinger et al., 2017. In the STIM condition, auditory stimuli (50 ms bursts of 1/f pink noise, inter-stimulus interval ≥ 2s) were played whenever the EEG signal in the C4 electrode (the “target channel”) crossed a default threshold of +30 µV (Figure 1, see Materials and Methods). Subjects were blind to the order of experimental nights and objective sleep parameters did not differ between STIM and SHAM nights except for the total time in bed (Supplementary Table 1).

**FIGURE 1.**
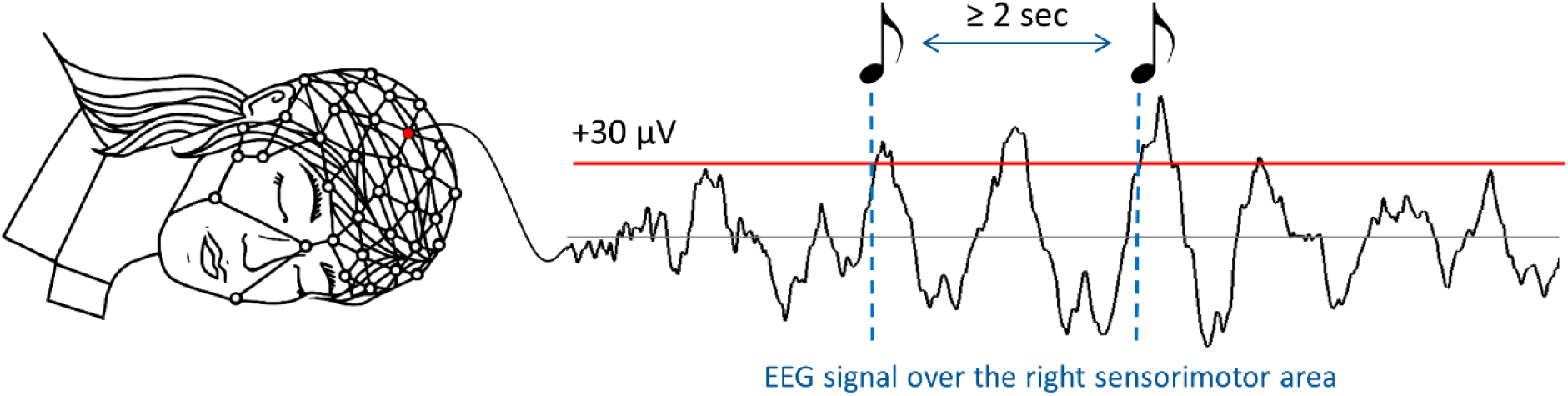
Closed-loop auditory stimulation protocol. During the STIM experimental nights, acoustic stimulation (50 ms bursts of 1/f pink noise, inter-stimuli interval ≥ 2s) was delivered whenever the EEG signal crossed a threshold of +30 µV (dashed lines) in the electrode C4 electrode located over the right sensorimotor area (red electrode on the high-density EEG cap).

### a. ERP and power analysis

We first explored the ERP and time-frequency representations (TFR) following the auditory stimuli presentation in the STIM condition and contrasted them to the SHAM condition, averaging across all channels. As shown in Figure 2, the presentation of auditory stimuli was followed by a pronounced ERP response, resembling a classical K-complex. The time-locked response contained all of the classical ERP components, including the P200, N350, N550 and P900 (Laurino et al., 2014). When comparing these ERPs with spontaneous slow waves in SHAM, we observed a significant change in the latency of the down-phase (t(1,8)=-4.77, p=0.001), occurring earlier in the STIM condition (0.55±0.03 s, M±STD) compared to the SHAM condition (0.72±0.11 s). A change in latency was also observed for the up-phase (t(1,8)=-4.89, p=0.001): again, the up-phase occurred earlier in the STIM (1.1±0.09 s) than in the SHAM condition (1.51±0.22 s). Not only the latency of down- and up-phase was shortened, but also the transition between down- and up-phase occurred faster (t(1,8)=-3.14, p=0.014), with a mean inter-peak latency of 0.53±0.72 s in the STIM condition and 0.76±0.21 s in the SHAM condition.

**FIGURE 2.**
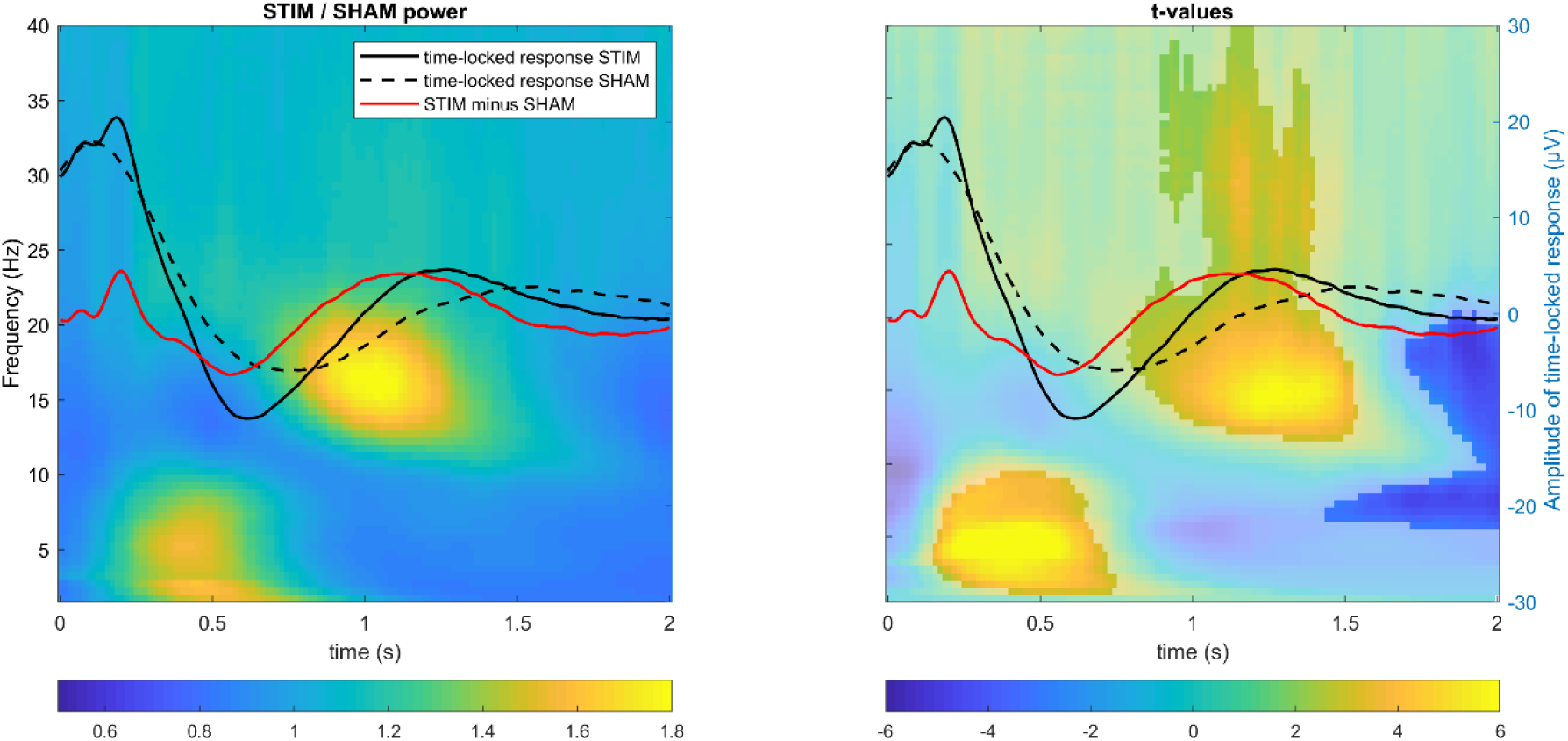
Total power changes following the stimulus during auditory closed-loop stimulation. Superimposed grand-averaged ERP waveforms (in C4 target channel) for STIM (solid line) and SHAM (dashed line) conditions in an overlay, with (*left*) time-frequency power plots (relative change in power between STIM and SHAM collapsed across all channels) and (*right*) time-frequency t-value plots (shaded area indicates non-significant difference between conditions, cluster corrected two-sided p<.05). Note that the stimulus presentation was associated with a power increase in the theta band during the transition to the down phase of slow waves, followed by a power increase in fast sigma and gamma bands during the up-phase of slow waves.

In the next step, we assessed total power differences between STIM and SHAM conditions across all frequencies from 0.75 to 40 Hz in the 2 seconds after stimulus onset using cluster-based permutation tests (Figure 2, TFR plots, average of all channels). This analysis showed that the activity in the delta (∼0.5–2 Hz) and theta range (∼3–10 Hz, prior to the slow-wave down-phase) was enhanced in STIM compared to SHAM by 48±35% and formed a cluster (p=0.02, d=1.31). Similar to previous findings, our TFR results have shown that up-phases of slow waves were associated with a robust increase of 44±26% in fast sigma power (p<0.001, d=0.77), with a center frequency of ∼16 Hz and extending to higher frequencies (slow gamma, ∼30 Hz). Interestingly, at the end of the 2-s analysis window, there was also a suppression of activity in the sigma band following the enhancement (p=0.03, d=-0.38; mean decrease - 13±6%), which is in line with previous findings on refractory periods of spindle occurrence (Antony et al., 2018).

To determine the spatial distribution of the observed effect, we characterized stimulation-related topographical changes in different frequency bands. A significant power enhancement within the delta band (0.75-3 Hz) was observed in frontocentral areas during the first second after stimulus onset (p=0.02, d=1.05, mean increase 41±34%). The widespread stimulation-related theta (4-8 Hz; p=0.003, d=0.98, mean increase 40±19%) and sigma (13-17 Hz; p<0.001, d=0.81, mean increase 51±34%) power enhancement also involved frontal and central areas. The time-resolved topographies in Figure 3 show that in both frequency bands the power increase arises as a local cluster of central channels, which spreads over frontal and parietal areas and lasts the longest over frontal areas. The cluster of power enhancement was distributed differently for theta and sigma components: in the theta band, the focus was centered around the vertex while in the sigma band the focus was in posterior channels over the parietal lobes.

**FIGURE 3.**
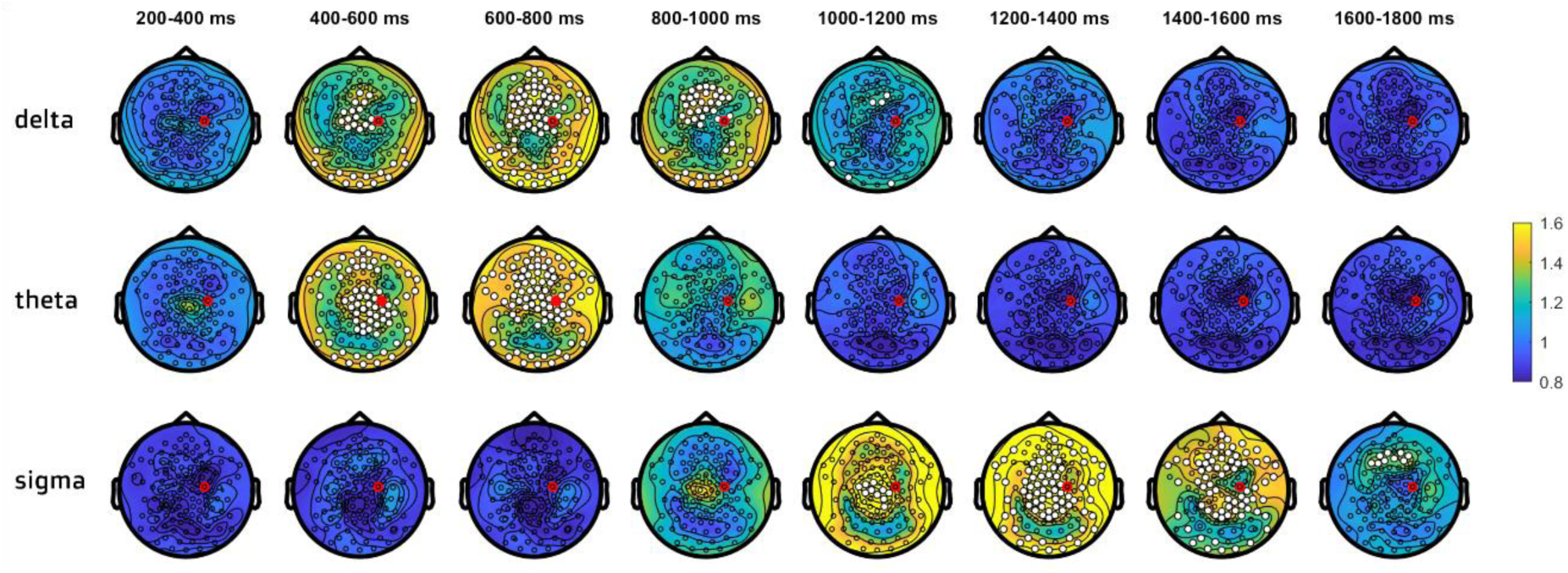
Topographical distribution of delta, theta and sigma power changes between STIM and SHAM conditions. Power value ratios (STIM/SHAM) were averaged in 200 ms time windows, starting from 200 ms after stimulus onset (start of the stimulation effect in the delta-theta range according to Figure 2). White dots indicate significant changes, cluster corrected two-sided p<.05. The red dot indicates the target channel C4.

In summary, this analysis shows a generalized enhancement of power in delta, theta and sigma bands following closed-loop acoustic stimulation, distributed predominantly in frontal and central areas.

### b. Cross-frequency coupling analysis

To characterize delta-theta and delta-sigma coupling in STIM and SHAM conditions, we compared the preferred delta phase and modulation index (MI) in both conditions in 9 representative channels across the cortex (F3, Fz, F4, C3, Cz, C4, P3, Pz, P4).

#### i. Coupling strength

To determine if delta-theta and delta-sigma coupling strength was affected by up-phase stimulation, we computed the MI at each channel in both conditions. The MI provides information only about the degree to which the amplitude of theta or sigma is modulated by the phase of delta oscillations. We normalized the coupling strength using a permutation-based reshuffling approach, resulting in a z-scored coupling strength (zMI), independent from differences in absolute power between conditions, channels, and individuals.

First, we separately assessed the coupling strength in STIM and SHAM conditions in the 9 representative channels. The coupling strength between delta and theta was significantly greater than zero both in STIM (all t(8) >5.32, p_adj_ < 0.001, FDR corrected) and SHAM (all t(8) >5.66, p_adj_ <0.001). The coupling strength between delta and sigma was also significantly greater than zero in STIM (all t(8) >2.97, p_adj_ < 0.02) and in SHAM (all t(8) >2.23, p_adj_ <0.05). The effect sizes were slightly larger in frontal channels for theta coupling (STIM, frontal channels: d=3.17, central: d=2.45, parietal: d=2.12) and in central channels for sigma coupling (STIM, frontal channels: d=1.14, central: d=1.89, parietal: d=1.07). Similar patterns of interregional differences were observed for the SHAM condition. These results were confirmed later by the topographical distribution of zMI (Figure 5).

**FIGURE 4.**
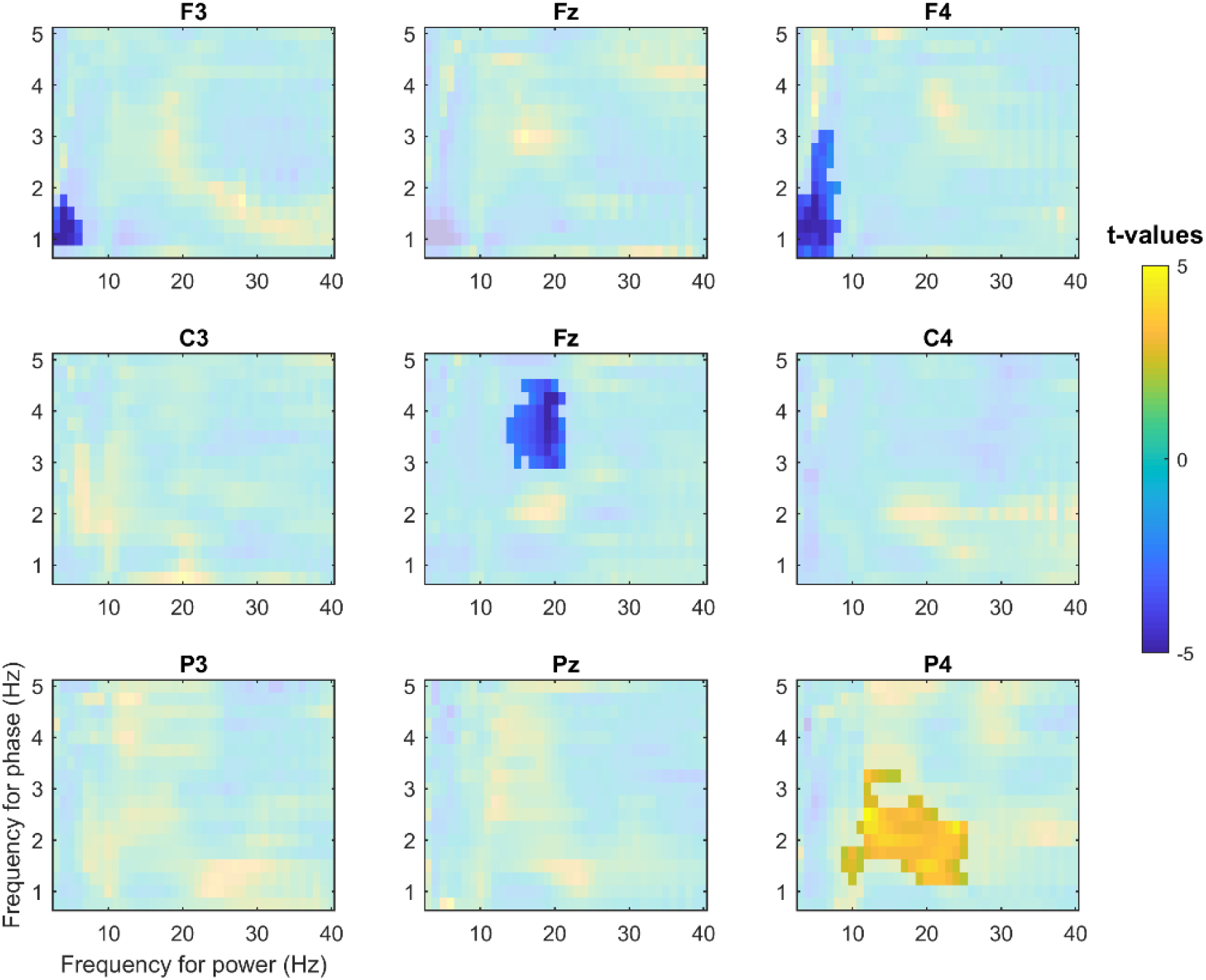
The modulation index as a function of amplitude (3 to 40 Hz) and phase (0.75 to 5 Hz) for nine electrodes. Non-shaded areas indicate statistically significant changes in cross-frequency coupling between STIM and SHAM conditions (cluster corrected two-sided p<.05). Reduction in coupling strength for delta and theta was observed in frontal channels F3 and F4 while coupling strength for delta and sigma bands was increased in the parietal electrode P4, located posterior to the target channel C4.

**FIGURE 5.**
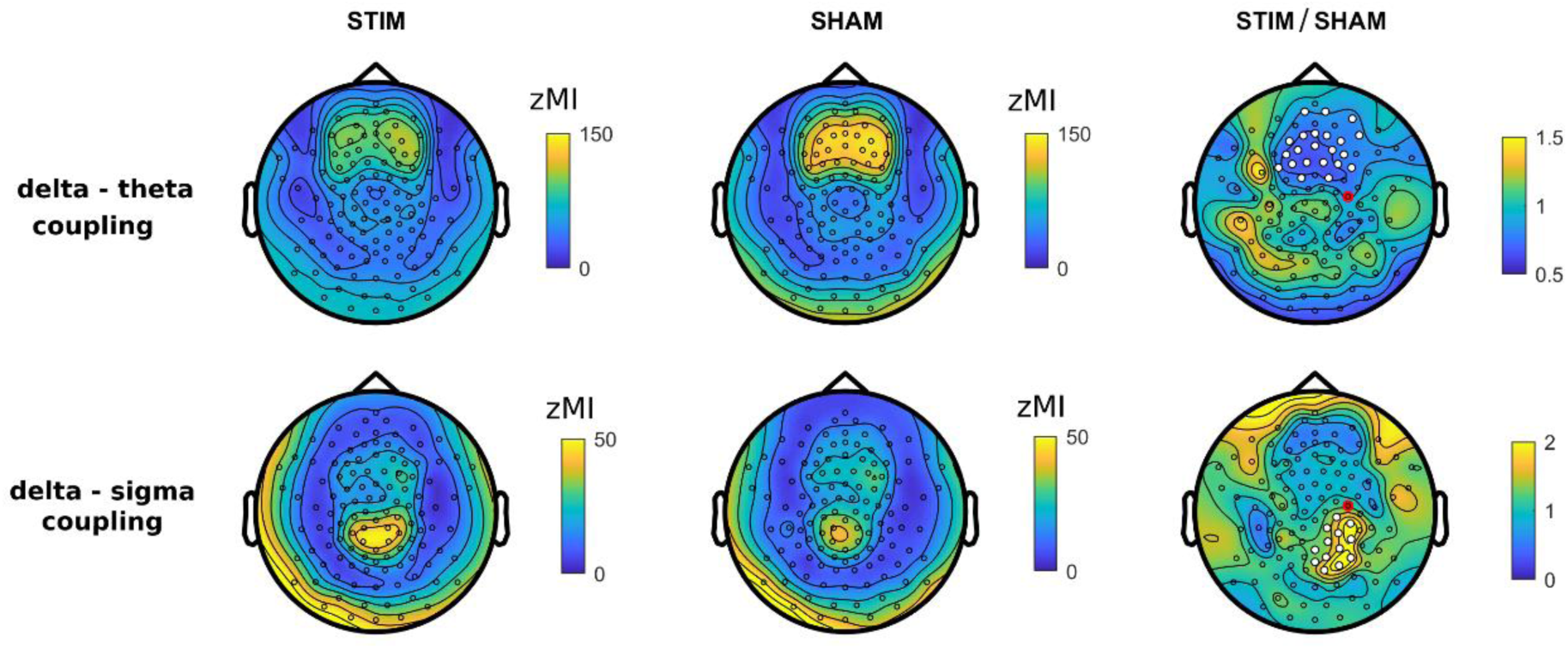
Topographical distribution of coupling strength. Delta-theta and delta-sigma coupling (modulation index) topography for STIM (*left*) and SHAM (*middle*) conditions and the ratio of STIM to SHAM (*right*). White dots indicate significant changes (cluster corrected two-sided p<.05), the red dot indicates the target channel C4.

Comodulograms in Figure 4 demonstrated region- and frequency-specific effects of the stimulation on coupling strength. A significant decrease in delta-theta coupling was observed in frontal channels F3 (p=0.004, d=-1.62) and F4 (p=0.01, d=-1.45) in the STIM condition as compared to SHAM. In the STIM condition, coupling strength for delta and sigma bands was increased in the parietal electrode P4 (p=0.004, d=1.2), located posterior to the target channel C4. Interestingly, in the vertex region, there was a significant decrease in coupling strength between the upper delta (∼3-4 Hz) and sigma (Cz, p<0.001, d=-1.85) bands.

#### ii. Topographical distribution of MI changes

Topographical distributions of delta-theta and delta-sigma coupling in STIM and SHAM conditions are shown in Figure 5. Delta-theta coupling was overall stronger in frontal sites both in STIM and SHAM conditions. For delta-sigma coupling, the focus was located over parietal sites. These results are in agreement with interregional differences in the effect sizes observed in 9 representative channels above.

We then compared MI topographies for the two conditions. The up-phase stimulation caused a significant decrease of delta-theta coupling by 31±18% in frontal channels (p=0.015, d=-1.64), the area of the strongest default coupling both in STIM and SHAM conditions. For delta-sigma coupling, we observed a significant increase by 124±105% over the right parietal area, located directly posterior to the target channel (p=0.01, d=1.15).

#### iii. Contribution of the phase-locked response to changes in delta-theta coupling

The overlay of TFR with ERP responses in Figure 2 indicates that the theta increase observed prior to the down-phase in STIM might contain non-phase-locked power changes (induced responses), as well as phase-locked power changes (evoked responses, including P200 and N350 components of the ERP). The majority of the theta changes that are observed 500 ms after stimulation are non-phase-locked, as can be seen on the induced power TFR in Figure S1 (for more details on phase-locked and non-phase-locked power analysis see Frequency analysis of the Methods section). A similar assumption can be made about activity in the delta range, containing both ongoing non-phase-locked activity overlaid with phase-locked stimulation-related components N550 and P900.

Along these lines, the observed decrease in coupling between delta and theta in frontal regions might be explained by an altered relationship between these two frequency bands due to the presence of evoked delta and theta responses. To test if the phase-locked activity in delta or theta bands is contributing to the observed decrease in coupling, in addition to zMI computed using total power (1), we calculated zMI using only: (2) broadband induced power (by subtracting the broadband ERP), (3) delta induced power (by subtracting the delta-filtered ERP) or (4) theta induced activity (by subtracting the theta-filtered ERP). Accordingly, zMI computed from total power includes both evoked delta and theta, whereas zMI computed from induced broadband power would not contain either of them.

A three-way ANOVA with factors *Condition* (STIM vs. SHAM), *EvokedDelta* (present or subtracted) *and EvokedTheta* (present or subtracted) yielded a significant interaction between *Condition* and *EvokedDelta* [*F*_(1,8)_=17.96, *p=*0.003, η^2^_*p*_=0.69]. The effect of *EvokedDelta* was significant for STIM [*F*_(1,8)_=32.76, *p*<0.001, η^2^_*p*_=0.8], but not for SHAM [*F*_(1,8)_=1.16, *p*>0.3, η^2^_*p*_=0.13]. Thus, only in the STIM condition, stronger coupling was observed when evoked delta activity was subtracted from the data (73.47±16.24 zMI) as compared to the coupling calculated including the evoked delta response (58.96±17.54 zMI). The analysis also revealed a significant main effect of *Condition* on coupling strength [*F*_(1,8)_=9.74, *p=*0.014, η^2^_*p*_=0.55], resulting from an overall weaker coupling in the STIM condition as compared to SHAM. The main effect of *EvokedDelta* was also significant [*F*_(1,8)_=18.50, *p=* 0.003, η^2^_*p*_=0.7]. The main effect of *EvokedTheta* was not significant, indicating that subtraction of the evoked theta did not affect zMI. No other significant interactions between factors were observed. Taken together, the ANOVA analysis demonstrated that the stimulation-related phase-locked delta could mediate the frontal decrease of cross-frequency coupling observed after up-phase stimulation.

### c. Delta phase preference

Theta and sigma oscillations were preferentially expressed during different phases of delta waves. Theta activity was maximal in the transition to the down phase of delta waves both for STIM and SHAM (average phase across 9 electrodes, averaged across subjects: 119±10° [circular mean ± SD] in STIM and 119±7° in SHAM; Figure S2). On the other hand, for the sigma band, the activity was preferentially expressed close to the up-phase of delta waves (−19±32° in STIM and -5±21° in SHAM; Figure S1). The phase distribution averaged across channels was non-uniform for theta and sigma power in both experimental conditions (Rayleigh test: all p<0.01).

The circular distributions in Figure S2 indicate the spatial extent of the preferred delta phase for theta and sigma frequencies. Because both theta and sigma maximal power values were scattered within half a cycle of the delta oscillation, we used a parametric two-way ANOVA with factors *Location* (frontal, central, posterior) and *Condition* (STIM vs. SHAM). Significant topographical differences in the preferred delta phase were observed both for theta and sigma. The distribution of the preferred phase formed a frontoparietal gradient, with the power maximum occurring earlier in the cycle in parietal areas. Thus, the maximum of theta power was located closer to the down phase in frontal sites and closer to the up-to-down transition in parietal sites [main effect of *Location*, F_(1.56,12.21)_=46.34; p<0.001, η^2^_*p*_=0.85; preferred delta phase: 143±3° in frontal, 118±3° in central and 93±5° in parietal sites]. Similar topographical differences were observed for sigma: the maximum of sigma power was located slightly after the up-phase of delta waves in frontal regions and slightly before in parietal sites [main effect of *Location*, F_(1.48,11.86)_=22.02; p<0.001, η^2^_*p*_=0.73; preferred delta phase: 28±13° in frontal, - 13±8° in central and -30±5° in parietal sites]. There was no difference in the mean coupling direction between STIM and SHAM condition for the theta band [main effect of *Condition*, F_(1,8)_=1.61; p>0.2, η^2^_*p*_=0.17]. However, across all locations, the maximum of sigma power occurred earlier in STIM (−12±7°) compared to SHAM (2±9°), as indicated by the significant main effect of *Condition* [F_(1,8)_=8.29; p=0.021, η^2^_*p*_=0.51]. No significant interaction between the factors *Condition* and *Location* was observed. In sum, acoustic stimulation significantly affected the preferred delta phase for sigma, but not for theta. In addition, we observed changes in phase preference across 9 representative electrodes.

## 3. Discussion

Although the majority of closed-loop auditory stimulation studies are still conducted in lab environments, recently, the first systems for unsupervised closed-loop enhancement of slow-wave activity have been developed (e.g., Debellemaniere et al. 2018; Ferster et al. 2019) with the goal to boost memory consolidation for prolonged periods and in clinical populations. Given the importance of local sleep characteristics for use-dependent neuroplasticity and considering the translational trend in the auditory closed-loop stimulation field, it is important to investigate whether slow waves, theta and sigma oscillations and their interaction can be manipulated selectively in a particular brain region. Here, we demonstrate that while closed-loop auditory stimulation globally enhances power in delta, theta and sigma bands, changes in cross-frequency coupling between these oscillations are expressed more locally. In particular, a significant increase in coupling between slow waves and sigma power was observed over the right parietal area, located directly posterior to the target electrode. This demonstrates that a local modulation of the coupling between slow waves and sigma band activity by closed-loop auditory stimulation could be used to manipulate sleep-dependent memory formation within the brain network of interest.

### Similarities between time-frequency profiles of stimulation-related responses and endogenous slow waves

By applying auditory stimulation with an inter-stimulus interval of ≥ 2 s, we had a sufficiently long time window to perform a detailed analysis of the ERP responses and associated spectral changes. Evoked ERP responses contained all the time-locked spectral components typical for stimulation-evoked K-complexes (P200, N350, N550, and P900) (Riedner et al., 2011). A time-resolved topographical analysis of the stimulation-related delta changes demonstrated a similar smooth transition between the near-vertex N350-like distribution and the frontocentral N550-like distribution within the first second after stimulation as found previously (Nicholas et al., 2002). K-complexes and the recently proposed type I slow waves (Siclari et al., 2014) are thought to be generated in sensorimotor areas and the medial parietal cortex and involve predominantly frontocentral regions (Bernardi et al., 2018; Caporro et al., 2012; Riedner et al., 2011; Siclari et al., 2014). Interestingly, stimulation-induced delta changes observed in our study were also distributed mainly in central areas, indirectly suggesting recruitment of the same neural mechanisms of slow-wave generation. Thus, a plausible interpretation of the commonly found closed-loop stimulation effects (e.g., Ngo et al., 2013) is that by applying stimulation time-locked to a particular phase of slow waves, further synchronization of endogenous slow waves is promoted by means of the externally evoked K-complex. This conclusion is in agreement with the interpretation of closed-loop auditory stimulation effects initially proposed by Bellesi et al. 2014. However, we cannot exclude that the frontocentral predominance of the observed changes in delta power may be related to the location of our target electrode C4. Shifting the target electrode to e.g. the occipital cortex would be an interesting probe of this idea for future experiments.

Similar to other studies using auditory stimulation during sleep (e.g., Cairney et al., 2018; Cox et al., 2014; Lehmann et al., 2016; Leminen et al., 2017; Papalambros et al., 2017), evoked responses within the delta range were associated with non-phase-locked activity in other frequency bands, namely theta and sigma. Changes in theta and sigma power were broadly distributed over the scalp and there was no target-electrode-specific local effect. However, there was a clear temporal relationship between slow waves and power in the theta and sigma frequency range as previously described in the literature and found not only after stimulus presentation, but also during spontaneous slow waves (Andrillon et al., 2011; Cox et al., 2014b, 2018; Gonzalez et al., 2018; Helfrich et al., 2018; Klinzing et al., 2016; Mölle et al., 2011). Interestingly, a very similar time-frequency pattern was observed in a TMS study, where single pulses over the primary motor area boosted sigma and theta power and these changes had a similar temporal relationship with down-phases (tms-N400) and up-phases (tms-P1000) of the evoked slow waves (Bergmann et al., 2012). Thus, theta-sigma enhancement associated with evoked K-complex-like responses might be of particular interest for understanding mechanisms underlying closed-loop auditory stimulation effects, as it represents the general “default” mechanism of brain responses to external stimulation.

### Effects of closed-loop auditory stimulation on coupling between the delta band and faster oscillations

Most recently, it has been shown that not only individual properties of slow waves and spindles can be associated with benefits of sleep for recovery and memory formation, but also the interaction between them (Demanuele et al., 2017; Helfrich et al., 2018; Latchoumane et al., 2017; Niknazar et al., 2015). The relationships between slow waves and sigma are well described in the literature. It is widely accepted that cortical down-states via cortico-thalamic connections are able to trigger activity in the sigma range which in turn is associated with the up-phase of delta waves (Staresina et al., 2015). Notably, there are very few scalp-EEG studies describing the generative mechanism of theta oscillations in NREM sleep and their relationships with slow waves.

In our study, we investigated the cross-frequency coupling between waves in the delta frequency range and faster oscillations in theta and sigma bands. Overall, despite the absence of local effects in power changes, auditory stimulation clearly modulated the local interaction between different spectral components. A comparison of the two experimental conditions revealed a stimulation-related decrease in coupling strength between delta and theta in frontal areas. This coupling seems not to be a target-channel-specific effect of the stimulation on the delta-theta coupling because it overlaps with the area of the strongest default coupling both in STIM and SHAM conditions. Taking into account that the delta and theta power enhancement contains both phase-locked (visible on ERP) and non-phase locked components (visible on TFR) the decrease in coupling between delta and theta in frontal sites might be explained by the altered interaction of these two components of the stimulation-related response. Our subtraction analysis of the evoked and induced components of delta and theta responses on coupling strength revealed that the stimulation-related phase-locked delta component primarily mediates the frontal decrease of coupling. One possible explanation for this observation might be that the slow part of the evoked K-complex propagating frontally is interacting with ongoing theta oscillations. Intracranial recordings could shed light on the interaction between delta and theta in baseline sleep and during stimulus presentation.

In contrast to non-target-electrode-specific delta-theta coupling changes, we found a region-specific effect for coupling between delta and sigma activity. The effect was observed for the area close to the target electrode, with a tendency to spread to the back of the head. This spread might be due to the traveling of slow waves, which primarily propagate from front to back (Massimini et al., 2004). An explanation of the local nature of the coupling between delta and sigma activity after stimulation might be related to the fact that slow waves are often local and not phase-locked across different brain areas (Nir et al., 2011). Given that the auditory stimuli are consistently time-locked to the up-phase in a specific (target) region, the effect might be restricted to that area. Consequently, the stimulation-evoked sigma increase peaked simultaneously with the next up-phase approximately one second after stimulation in that same area.

We also found stimulation-related changes in the preferred phase of sigma oscillations. Following stimulation, the sigma power peak occurred slightly prior to the positive peak of the slow wave, whereas in the SHAM condition power peaked slightly after the positive peak. This is consistent with the observation that external stimulation evokes fast spindles, which are known to appear earlier in the delta cycle compared to slow spindles. Enhancement in the sigma band was also distributed in parietal areas, confirming previous findings on the topography of fast spindles (Cox et al., 2017; Mölle et al., 2011).

In summary, we observed two distinct effects of closed-loop auditory stimulation on cross-frequency coupling for two frequency bands involved in memory consolidation during sleep. In many previous studies, transcranial direct current stimulation was used to demonstrate the relevance of the delta-sigma coupling for the overnight memory formation (for example, Marshall et al. 2006; Westerberg et al. 2015; Ladenbauer et al. 2016). The global sigma boost observed one second after auditory stimulation could provide the optimal time-window for memory replay, with content specificity encoded in the gamma band. In a recent iEEG study, it was shown that the sequences of population firing peaks across widespread cortical regions in the gamma band recorded during wake are re-emerging during NREM sleep. Occurrences of these “motifs” were coupled to down-to-up slow-wave transition, spindles and hippocampal sharp-wave ripples (Jiang et al., 2017). Along these lines, closed-loop auditory stimulation targeting slow waves in a particular brain region could facilitate the memory replay with particular content by promoting better local coordination between delta, sigma, and nested gamma. In future studies, it will be important to clarify if closed-loop auditory stimulation might improve region-specific memory consolidation.

### Conclusion

Our data suggest that closed-loop auditory stimulation might provide a minimally invasive way to modulate the relationships between the main sleep rhythms in a target region while not altering sleep architecture. This opens the way to a large range of future implementations of closed-loop auditory stimulation in different clinical populations. For instance, assessment and local manipulation of cross-frequency coupling during NREM sleep could shed light on the pathophysiological mechanisms underlying sleep-related learning problems observed in clinical populations such as schizophrenia (Chan et al., 2017; Ferrarelli, 2015) or epilepsy (Höller and Trinka, 2015).

## 4. Materials and Methods

### Participants

Nine right-handed subjects (23±1.3 y.o., 4 females) participated in the study. Participants took notes on their daily activities, sleep and wake time in a sleep diary and were actigraphically monitored (Actiwatch Type AWL from Cambridge Neurotechnology, CamNtech, Cambridge, UK or Geneactiv) for the 7 days (range from 4–8 days) preceding the experiment. Participants did not take any medication at the time of the experiment and were required to refrain from alcohol and caffeine 24 h prior to each experimental session. The sleep schedule in the laboratory was organized in accordance with individual sleep habits. Written informed consent was obtained prior to participation. The study was approved by the local ethics committee and performed according to the Declaration of Helsinki.

### High-density sleep electroencephalography

Full-night sleep was recorded by using high-density EEG (Electrical Geodesics Sensor Net for long-term monitoring, 128 channels, referenced to a vertex electrode, sampling frequency 500 Hz). Submental electromyographic and electrooculographic data were collected for visual sleep scoring. Two additional electrodes (gold, Grass Technologies, West Warwick, RI, USA) were attached to the earlobes, which served as reference electrodes for the online closed-loop auditory stimulation. After adjusting the net to the vertex and the mastoids, all electrodes were filled with an electrolyte gel to ensure the maintenance of good signal quality throughout the night. EEG electrode impedances were below 50 kΩ. Impedances were below 20 kΩ for the electrodes used for online closed-loop auditory stimulation: submental electrodes, electrodes on the earlobes, and the electrode used for slow-wave detection (C4).

### Experimental protocol

Subjects participated in two sessions carried out on different nights separated by 1 week: (i) non-stimulation (SHAM); (ii) closed-loop auditory stimulation in the up-phase (up STIM) of real-time detected slow waves. The order of the STIM and SHAM sessions was randomized and counterbalanced across subjects. In the STIM condition, closed-loop auditory stimulation (STIM) was applied precisely time-locked to the up phase of sleep slow waves detected in the C4 electrode. Real-time closed-loop slow-wave detection during N2 and N3 was performed as described in Fattinger et al., 2017. In the STIM condition, auditory stimuli (50 ms bursts of 1/f pink noise, 50 dB with a rising/fall time of 5 ms, inter-stimulus interval ≥ 2s) were played whenever the EEG signal crossed a default threshold of +30 μV. Stimulation was immediately stopped in case of an arousal and if the amplitude of the continuously monitored EMG exceeded a given threshold set by the experimenter.

### Data analysis and statistics

EEG data were analyzed using Matlab R2017b, using custom-written scripts and the FieldTrip toolbox (http://fieldtrip.fcdonders.nl/, Oostenveld et al. 2010).

### Sleep scoring

For sleep scoring, the EEG data were bandpass filtered (0.5–50 Hz), re-referenced to the earlobes and downsampled to 128 Hz. The EEG was visually scored by two trained individuals for sleep stages Wake, N1, N2, N3 and REM sleep at frontal, central and occipital electrodes (20 s epochs) based on American Academy of Sleep Medicine standard criteria (Iber et al., 2007).

### EEG preprocessing

Data were bandpass filtered (0.5–45 Hz, mirror padding: 3 s) and preprocessed in 10 s epochs around the stimulus onset, with a 6 s prestimulus interval. In the SHAM condition, where no stimuli were applied, slow-wave up-phase detection was performed offline. Epoched data were downsampled to 128 Hz to speed up processing. Channels located on the earlobes and on the face below the front were excluded from further analysis. After visual inspection, noisy channels and trials containing strong movement artifacts were removed (on average, no more than 5 sensors and 5.8 trials were removed per session). Removed channels’ values were interpolated using spherical interpolation. The cleaned data were re-referenced to an average value across all 118 channels. Next, for a consistent phase relationship between the auditory stimuli and the slow waves, we selected only the trials where the stimulus was applied during the rising part of the positive half-wave. For this, we filtered the signal between 0.5-2 Hz and we calculated the instantaneous phase of each slow wave at stimulus onset after applying a Hilbert transformation. Approximately 60% of the total number of trials were selected for further analysis. We used a 6 s interval prior to the stimulus onset for baseline correction.

### Frequency analysis

To identify global spectral responses evoked by acoustic stimulation, we performed time-frequency analyses separately for total and induced power changes. Total power contains both evoked power (time-locked and phase-locked response to stimulus presentation) and induced power (time-locked but non-phase-locked response). To compute the total power values in STIM and SHAM conditions, single-trial time-frequency representations (TFRs) of power were calculated by FFT with a Hanning taper applied through an adaptive time window of 5 cycles for each frequency between 0.75–40 Hz (ΔT = 3/f for 0.75–3 Hz and ΔT = 4/f for 3–40 Hz) in steps of 0.008 s. Power estimates in the time-frequency domain were averaged across all EEG channels and trials. Through visual inspection of the TFR plot we identified three main spectral contributions in delta (0.75–3 Hz), theta (4–8 Hz) and sigma (13–17 Hz) bands and described their spatiotemporal evolution through topographies. To obtain induced power values, the non-phase-locked activity was calculated by subtracting the ERP from each trial (separately for each condition) and then performing time-frequency analysis on the residual time series.

### Cross-frequency coupling

Given the presence of distinct spectral components modulated by the acoustic stimulation, we quantified their relation in terms of cross-frequency coupling strength and preferred coupling phase.

Modulation index (MI) (Canolty et al., 2006) indicating the association between phases in the 0.75-5 Hz (delta) range and power in the 3-40 Hz band was calculated for each 2-s epoch using fieldtrip software (Oostenveld et al., 2010). This method is based on the following: under the null-hypothesis of no systematic cross-frequency coupling, the observed amplitude distribution of fast frequency over phase bins of slow frequency does not deviate from the uniform distribution and MI is zero. If the amplitude distribution over phase bins significantly deviates from uniform, the MI will deviate substantially from zero, indicating cross-frequency coupling. For this analysis, we used 18 bins following Tort et al., 2010. Importantly, coupling strength as assessed by MI depends on the signal-to-noise ratio, which might be confounded by power differences in STIM and SHAM conditions. Therefore, for every subject and channel, we constructed a null distribution of MI by repeatedly (*n* = 200) shuffling the delta phase timeseries with respect to the fast frequency amplitude timeseries, and recalculating the MI for each iteration. This null distribution was used to Z-transform the empirical MI, resulting in a zMI that is independent of power differences.

The preferred phase analysis was performed in the time-window of 2 s after stimulus onset. After applying a Hanning window (parameters similar to the frequency analysis described above) and obtaining complex Fourier coefficients for the low (delta) and the high (theta or sigma) frequency bands, we computed the absolute amplitude of the high frequency signal in each epoch and sorted the resulting values according to the instantaneous phase of delta oscillation in 180 bins.

### Statistics

To correct for multiple comparisons when examining statistical differences between two conditions, we used a nonparametric clustering procedure (Maris, 2012; Maris and Oostenveld, 2007). First, independent samples t-tests were computed (2-tailed, p<0.05) for the difference between STIM and SHAM conditions for all tiles in the time-frequency domain. Next, significant neighboring time-frequency tiles were clustered if they showed the same direction of effect. To assess statistical significance of each cluster, a cluster-level test statistic was calculated by computing the sum of all t-values in the cluster. The significance of each cluster-level statistic was estimated by comparing the cluster-level test statistic to a reference permutation distribution derived from the data. The reference distribution was obtained by randomly permuting the data 5000 times. The cluster *p*-value was estimated as the proportion of the elements in the reference distribution exceeding the cluster-level test statistic. An analogous procedure was used for the statistical comparison of zMI frequency-frequency plots. Statistical significance of topographical changes was tested in a similar manner: clustering was applied by pooling together neighboring electrodes.

Two-way repeated measures analysis of variance (ANOVA) was used to assess the influence of factors *Location* (frontal, central, posterior) and *Condition* (STIM vs. SHAM) on the preferred delta phase separately for theta and sigma bands. A three-ways repeated measures ANOVA with factors *Condition* (STIM vs. SHAM), *EvokedDelta* (present or subtracted) *and EvokedTheta* (present or subtracted) was used to assess the contribution of time-locked components in the coupling of delta and theta. We used the Greenhouse–Geisser correction to estimate the *p-*values. The level of significance was set to *p* < 0.05.

The circular Rayleigh test was used to determine whether circular coupling phase distributions deviate from the uniform distribution. All statistical tests were two-sided.

## 5. Acknowledgements

This work was supported by the Swiss National Science Foundation (grant number 320030_153387), the Clinical Research Priority Program (CRPP) “Sleep and Health” of the University of Zurich Switzerland and the HMZ Flagship Project SleepLoop of University Medicine Zurich Switzerland.

---

**Supplementary Table 1.** Sleep architecture. Sleep parameters were calculated from the visual scoring of sleep stages. Sleep efficiency was calculated by dividing total sleep time with total time in bed. Wake after sleep onset is expressed as the percentage of total time in bed. Sleep stages (NREM sleep, stage N1, stage N2, stage N3, and REM sleep) are expressed as a percentage of total sleep time. P-values result from paired Student’s t-tests. Bold letters indicate a significant difference (p < 0.05). Sleep architecture did not differ between the two nights except for total time in bed.

|  | STIM |  | SHAM |  | p-value |
| --- | --- | --- | --- | --- | --- |
|  | mean | SD | mean | SD |  |
| **Sleep architecture** |  |  |  |  |  |
| Total time in bed [min] | 451 | 21 | 468 | 12 | **0.03** |
| Total sleep time [min] | 410 | 35 | 430 | 22 | 0.08 |
| Sleep efficiency [%] | 91 | 5 | 92 | 3 | 0.63 |
| Sleep latency [min] | 24 | 19 | 18 | 9 | 0.31 |
| Wake after sleep onset [%] | 5 | 4 | 5 | 3 | 0.79 |
| NREM sleep [%] | 79 | 5 | 78 | 5 | 0.96 |
| Stage N1 [%] | 5 | 2 | 5 | 3 | 0.81 |
| Stage N2 [%] | 51 | 7 | 51 | 9 | 0.99 |
| Stage N3 [%] | 23 | 8 | 23 | 7 | 0.75 |
| REM sleep [%] | 21 | 5 | 21 | 6 | 0.96 |

**FIGURE S1.**
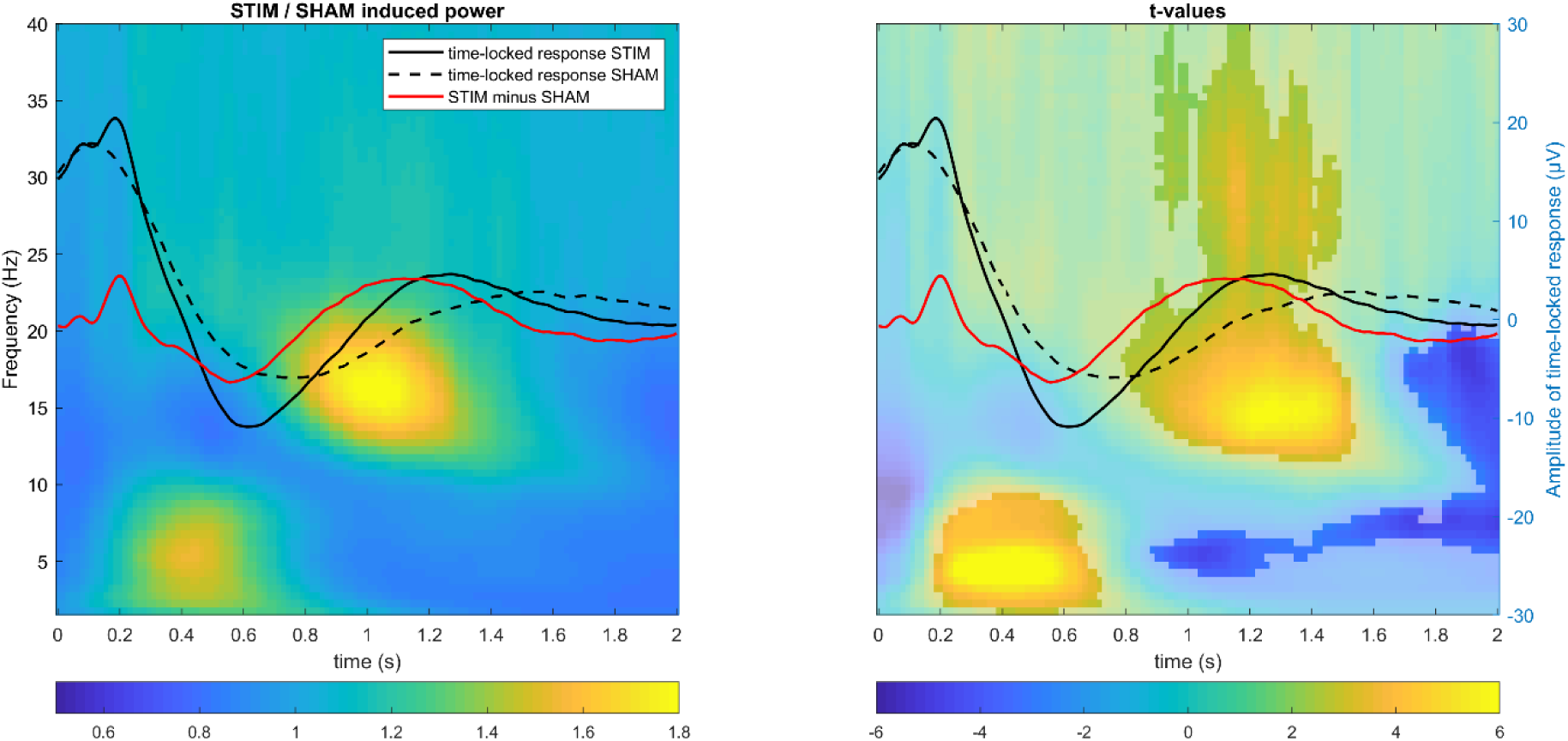
Induced power changes following the stimulus during auditory closed-loop stimulation. Superimposed grand-averaged ERP waveforms (in C4 target channel) for STIM (solid line) and SHAM (dashed line) conditions overlaid with (*left*) time-frequency power plot (relative change in power between STIM and SHAM collapsed across all channels) and (*right*) time-frequency t-values plot (shading indicates extent of significance, cluster corrected two-sided p<.05). Note that the induced power increase in theta and sigma bands is similar to the total power increase in Fig 2.

**FIGURE S2.**
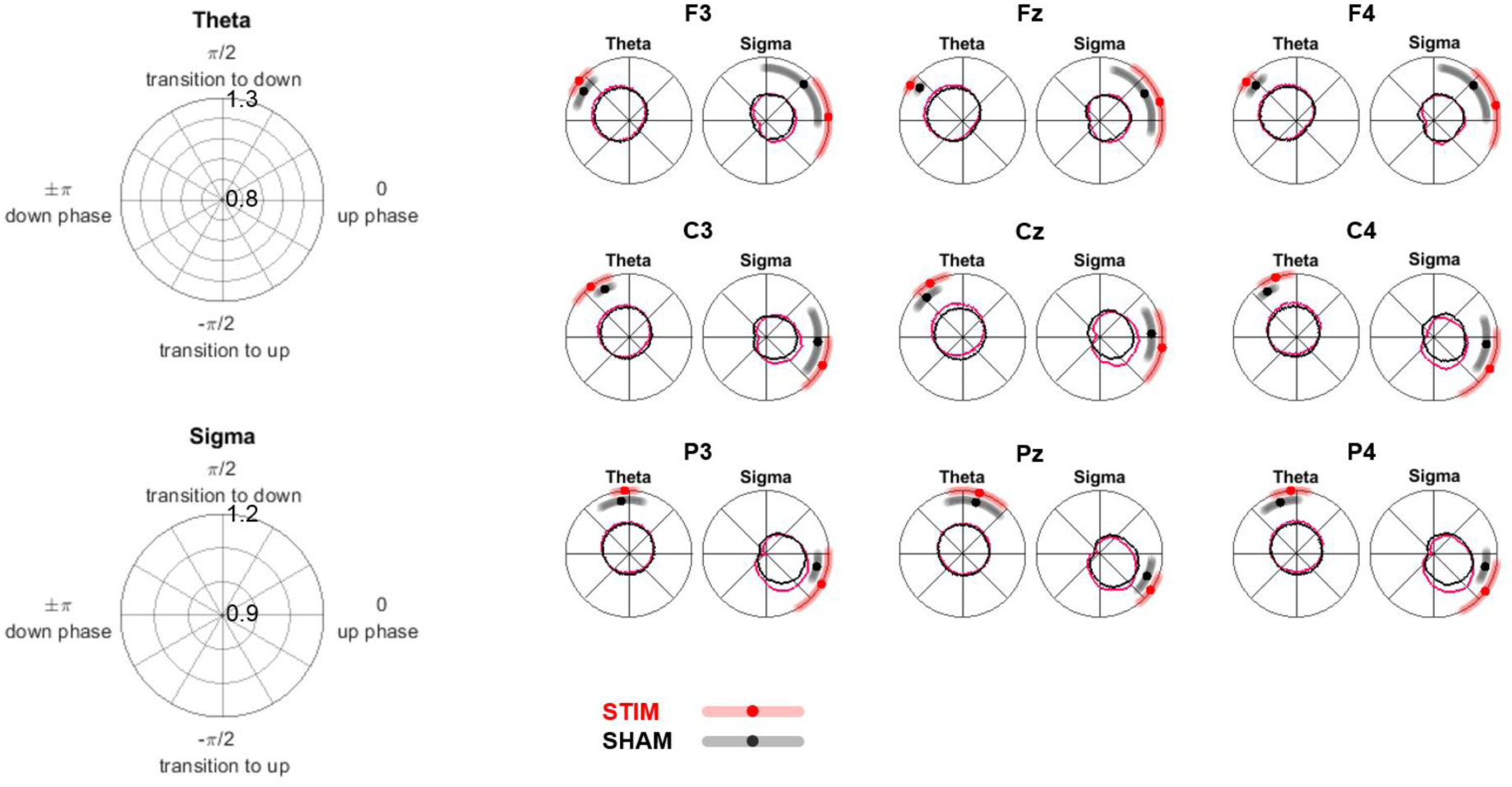
Delta-sigma coupling phases across 9 representative channels. Dots and shading on polar plots display the mean preferred phase of delta waves (±STD) for 9 subjects. The radius of the circle inside the polar plot illustrates the dependence of the mean normalized amplitude of fast oscillation (theta or sigma) on the delta phase.

